# No exploitation of temporal predictive context during visual search

**DOI:** 10.1101/2020.08.25.265975

**Authors:** Floortje G. Bouwkamp, Floris P. de Lange, Eelke Spaak

## Abstract

The human visual system can rapidly extract regularities from our visual environment, generating predictive context. It has been shown that spatial predictive context can be used during visual search. We set out to see whether observers can additionally exploit temporal predictive context, using an extended version of a contextual cueing paradigm. Though we replicated the contextual cueing effect, repeating search scenes in a structured order versus a random order yielded no additional behavioural benefit. This was true both for participants who were sensitive to spatial predictive context, and for those who were not. We argue that spatial predictive context during visual search is more readily learned and subsequently exploited than temporal predictive context, potentially rendering the latter redundant. In conclusion, unlike spatial context, temporal context is not automatically extracted and used during visual search.

## Introduction

From the moment we wake up, we are continuously confronted with visual input. Selective attention helps us to navigate this rich visual environment, by enabling us to process what is important for the task at hand, and to ignore the rest. For instance, a bike ride to work requires us to attend the road, the signs and traffic, without letting our eyes wander to the bright blue sky. Humans are incredibly good at focusing attention to select objects from cluttered scenes. An important factor that contributes to this ability is that our visual system makes use of predictable structure in our environment (1,2). Our world might be very complex, but it is also very stable: the road is always beneath us and signs are usually found at the roadside. Co-variation between objects and their environments creates a predictive context that can be exploited by the visual system to guide selective attention more efficiently. There is indeed ample evidence that contextual expectations can help us identify objects in cluttered scenes (1–4). Our visual system appears to be geared to automatically and implicitly extract regularities from our experiences (5,6) and use these regularities, or *predictive context*, to process the world more efficiently (7).

However, we navigate through the world not only in space, but also in time. On the aforementioned route to work, we use *spatial* context to locate traffic lights faster, but we also know, for example, that this particular roundabout will follow that specific turn. In other words, in everyday life, we might additionally make use of *temporal* context.

One way to investigate *spatial* predictive context is by using the well-established contextual cueing effect (7,8). A contextual cueing experiment consists of a visual search task during which scenes, typically one T-shaped target embedded amongst similar looking distractors, are repeated. Though unaware of these repetitions, people become markedly faster in finding the target in these repeated scenes. Consensus is that contextual cueing leads to a more efficient guidance of spatial selective attention through the search displays (8–11).

To investigate *temporal* predictive context, single items are generally shown in a specific order, i.e. as a pair, triplets, or longer sequences. Due to this order, one item becomes predictive of the following item(s). These items can be syllables (12), shapes (6), or objects (13–15). Similarly to the spatial regularities in contextual cueing, this temporal regularity typically goes unnoticed by participants.

It is unclear whether observers exploit temporal predictive context, in addition to spatial regularities, when locating an item in a cluttered scene. We therefore set out to see whether the contextual cueing effect extends to the temporal domain. Can observers learn from both spatially predictive and temporally predictive context in visual search?

To answer this question, we exposed participants to various types of predictive context within a visual search task. As in classical contextual cueing, we repeated displays many times throughout the experiment, and intermixed them with completely novel displays. We therefore anticipated a behavioural benefit for the repeated scenes, replicating the classic effect of spatial predictive context. As a novel manipulation, we presented repeated displays either in a random or in a predictable order. Therefore, if observers can additionally learn from temporal predictive context, we would expect a search time benefit when search scenes are repeated in a consistent order, compared to when they are repeated in a random order.

To preview our main results: we find a strong benefit of repeating scenes, but no additional advantage of presenting them in a consistent order. We conclude that, in the case of visual search, spatial predictive context might be dominant, rendering temporal predictive context redundant. This finding increases our understanding of how the visual system exploits previous experience to process the complex world more efficiently.

## Materials and methods

### Data availability

All data and code used for stimulus presentation and analysis are released publicly under an open license at the Donders Repository [http://doi.org/10.34973/hrcx-8w07] and will be available at publication.

### Participants

We recruited 40 healthy participants with normal or corrected-to-normal vision. Two participants were excluded as they missed too many trials (proportion late trials 38.54% and 29.98%, both scores fall below 25th Percentile minus 1.5 x Interquartile range), resulting in a final sample of 38 participants (27 women, age 19-36 years, *M*=24.58, *SD*=3.97). This sample size was chosen to yield >80% power to detect differences with a medium effect size (d = 0.5) with a two-sided paired t-test at an alpha level of 0.05 (requiring n = 34), plus margin. The experiments were approved by the local ethics committee (CMO Arnhem-Nijmegen, The Netherlands) under the general ethics approval (CMO 2014/288, version 2.1) and were conducted in compliance with these guidelines. All participants gave written informed consent beforehand and were paid for their participation.

### Procedure

All stimuli measured 1.2° x 1.2° in size and were presented as black on a grey background. Participants were instructed to search for a T-shaped target stimulus amongst nine L-shaped distractors. Distractor L shapes had a 10% offset in the line junction to increase search difficulty (16) and were rotated a random multiple of 90°. The target T was tilted either to the left or to the right (90°), and the participant was instructed to indicate the orientation of the T with a left or right button press. Throughout the experiment, a fixation dot (outer white diameter 8 pixels; inner black diameter 4 pixels) was presented at the center of the screen. Participants were asked to fixate and not move their eyes. Each trial started with a 1 s fixation period, and search displays were presented for 2.5 s or until the response button press. After the button press, participants were informed whether their response was on time and correct or not by the outer part of the fixation dot turning green (correct), red (incorrect) or blue (too late) for 0.5 s (Figure 1A). A regular grid with steps of 1°, spanning from −9° to +9° horizontal and −6° to +6° vertical from the center of the screen was used to arrange stimuli on the screen. Random jitter of ± 0.4° was added to stimuli locations to prevent collinearities with other stimuli (8). To ensure displays were of equal difficulty, the target always appeared between 6° and 8° of eccentricity, and the mean distance between the target and all distractors was kept between 9° and 11°. Additionally, both local crowding (too many distractors surrounding the target) and a ‘standing out effect’ (too little distractors surrounding the target) were not allowed. To further equate difficulty between the Old conditions (see *Experimental conditions*), we counterbalanced displays sets between the Old-ro and Old-so conditions across participants.

**Figure 1.**
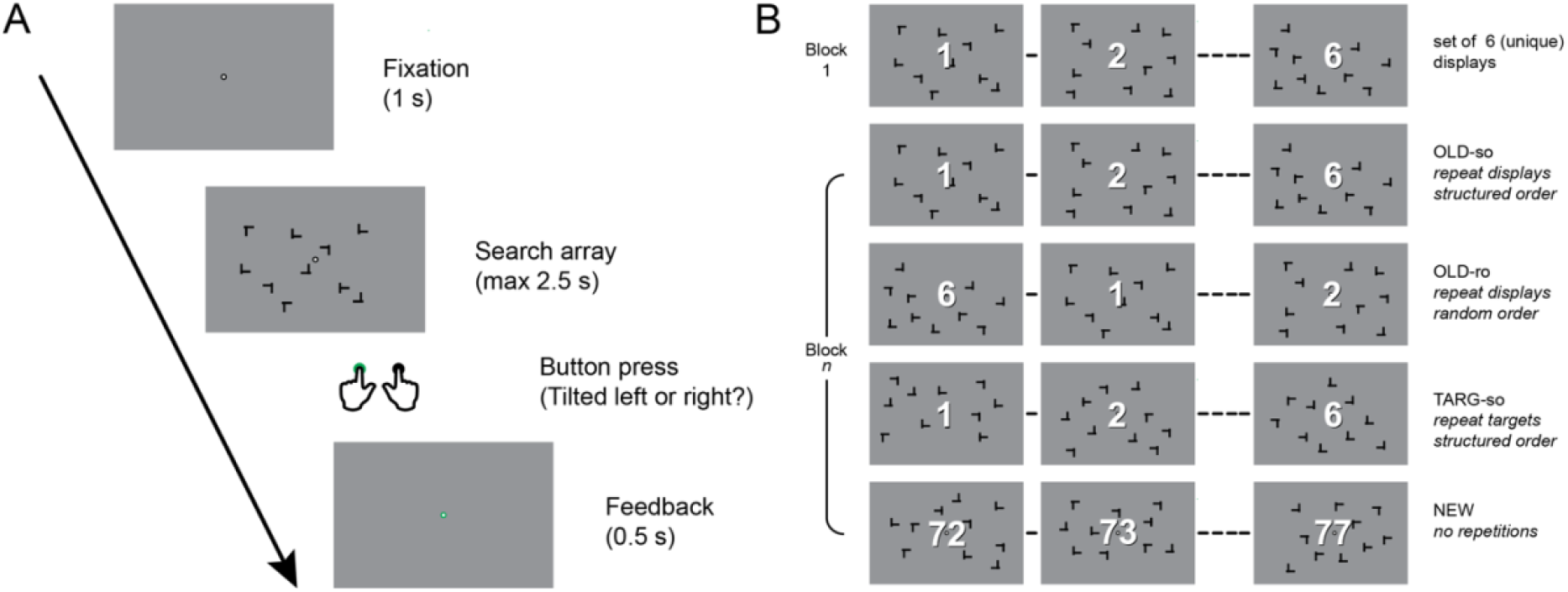
Search task and experimental design. (A) Schematic of an individual search trial. (B) The different conditions are defined by what is repeated on subsequent blocks (whole display or target location only) and in what order these are repeated (random or structured order). The same set of displays is shown here for illustration purposes only, in reality a unique set of displays was used for each condition.

After practicing the task for 72 trials, the main search task commenced, consisting of 36 blocks of 24 trials each. Participants were given feedback on their performance (% correct) and could take a short break every third block.

### Experimental conditions

For each condition, six unique displays were repeated in a specific manner. Both the location and rotation of all distractors as well as the location of the target were the same throughout the experiment for *Old* displays. The tilt of the target (leftward or rightward) was however randomized to prevent participants from learning the correct motor response for a given display. This way, the layout of the distractors was only of predictive value for the target location, generating *spatial predictive context*. The six displays were represented together as a sub-block, but the order within the sub-block however was randomized for the Old-random-order (Old-ro) condition. For Old-structured-order (Old-so) trials, this order was always the same. This predictable order added *temporal predictive context*. We also included a Target-structured-order (Targ-so) condition. Instead of the entire configuration, only the location of the target was repeated across blocks for Targ-so displays, and this repetition of target locations was again in the same order across blocks. Target location on the first trial therefore predicted the next target location, and so on, for all 6 Targ-so displays. All search blocks contained a sub-block of six ‘New’ trials, i.e., newly randomly generated search displays. Together this yielded four conditions, each containing six displays per sub-block. The order of these sub-blocks was randomly permuted within an experimental block, with the constraint that an immediate repetition of a condition was not allowed (i.e., the last condition of block n was not allowed to be the same as the first of block n+1).

After the main search task, we probed explicit knowledge of the presented spatial and temporal regularities. Before testing participants knowledge, we inquired about their subjective experience of recognition with the question: “Did you have the feeling that some of the search displays occurred multiple times over the course of the experiment?” and indicated their (unspeeded) answer (‘yes’ or ‘no’) with a button press. Next, they were probed about their confidence: “How sure are you about your answer to the previous question?” and answered with a button press (‘very sure’ or ‘not very sure’). We pooled responses across confidence levels, since 74 % (28/38) of participants indicated ‘not very sure’ to this confidence question. Subsequently, participants performed a recognition task. The 12 Old trials from the main experiment and 12 newly generated New trials were randomly intermixed and participants had to indicate whether a display was used during the main search task or not. Participants could take their time answering the question (no time outs) and did not receive feedback.

Finally, participants were probed on their explicit knowledge of the structured order conditions. We inquired about their subjective experience with the question “Some of the displays that were repeated were also always presented in the same order. Did you notice this?” and asked to answer with a button press (‘yes’ or ‘no’). Again they were asked to indicate their confidence in the previous answer. After this, participants performed another recognition task, this time on the order of two displays that were presented consecutively. We presented 20 trials of two displays each from the structured order sets (Old-so and Targ-so) and presented them either in the correct or incorrect order.

Participants were instructed to indicate if the presented order on a given trial was the same as during the main search task, or different. Only 4 participants (10%) reported having noticed a structured order of repetitions, and only half of them were confident in their answer. We therefore ignored both the subjective recognition rating and the confidence rating in the analyses of the data from this recognition task.

### Apparatus

Stimuli were presented on a BenQ XL2420T monitor, at a resolution of 1920 × 1080 pixels and a refresh rate of 120 Hz, using Matlab (The Mathworks, Inc., Natick, Massachussetts, United States) and custom-written scripts using the Psychophysics Toolbox. Participants were seated in a separate room with dimmed light, at a distance of 55 cm from the screen using a chinrest.

### Data analysis

Data analysis and visualization was done using R (17). Reaction time (RT) was our primary, and accuracy our secondary variable of interest. Statistical assessment of the RT data was done for the second half of the experiment, i.e. blocks 18–36 (selected a priori). Only RT data from correct trials was used. For the structured order conditions (Old-so and Targ-so), RT data for the first trial of each subblock was discarded, as no temporal predictions can be made for the first display of the set. Since RT distributions are typically heavily skewed, the RT data was log-transformed prior to any statistical analysis to improve normality.

To prevent participants from forming a more general prediction of target location per quadrant of the screen (18) we balanced target location per condition as much as possible across the quadrants of the screen, and regressed out any residual effect of quadrant frequency (QF) from the data per participant. For visualization purposes only, we smoothed the timecourses of both RT and accuracy across neighbouring blocks (i.e., the data plotted at block n represents the mean of blocks n−1, n, and n+1; Figure 2A). All plots were generated with the ggplot2 package for R.

**Figure 2.**
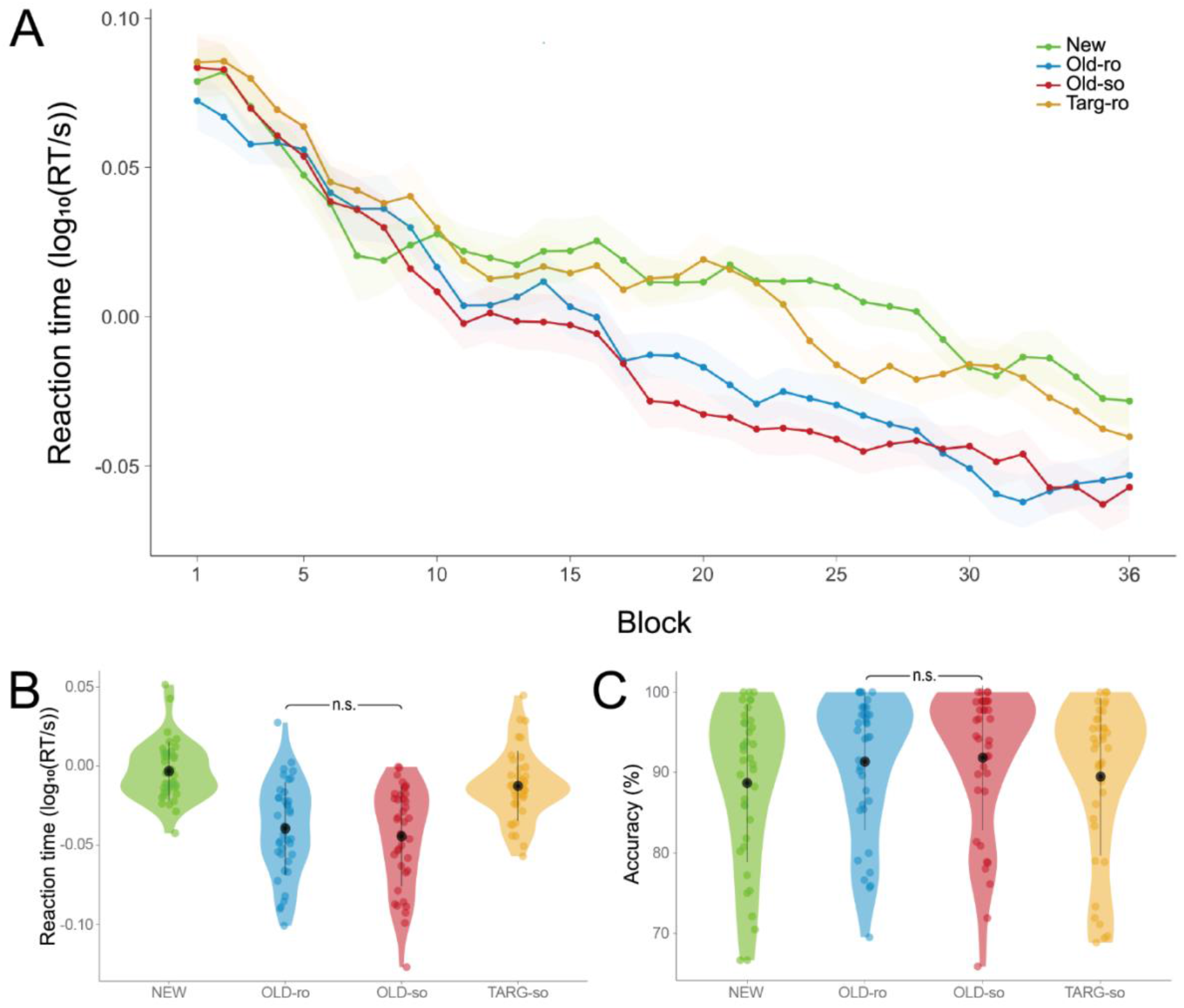
Results main search task. (A) Reaction time (smoothed) plotted over the timecourse of the experiment (shading indicates within-participant corrected standard error of the mean) (B) Reaction time and (C) Accuracy within the second half of the experiment (block 19-36). Colored dots are individual participants, the black dot reflects the mean, and the black bar indicates ±1 standard deviation.

We report raw RTs in the Results section, but all statistical tests were done on log-transformed and QF-corrected quantities. If the assumption of sphericity was violated, as indicated by a significant outcome of Mauchly’s test, we report the corrected p value and degrees of freedom. All Greenhouse-Geisser ε values were above 0.75, we therefore reported the more liberal Huyn-Feldt corrected values (19). Effect sizes were calculated using the effsize package for R. Specifically, the value of Cohen’s d is computed using the approach of Gibbons et al. (20) for paired samples, including a suggested correction of Borenstein (21).

In addition to frequentist statistics, we report the Bayes Factor. We applied a standardized Bayesian t-test, using the BayesFactor package for R. The Bayes factor quantifies the relative evidence for the alternative compared to the null hypothesis. The outcome is a ratio that quantifies how much more likely the data are under one hypothesis, than under the other. In our case, values >1 mean the data are more likely under the alternative hypothesis than under the null hypothesis, and conversely, values <1 mean the data are more likely under the null hypothesis than under the alternative hypothesis.

## Results

### Visual search task

Both search speed and accuracy improved over the time course of the experiment (Figure 2A), revealed by a main effect of time (first/second half, reaction time: F_1,37_ = 174.4, p = 1.4 × 10_−15_, accuracy: F_1,37_ = 16.7, p = 2.3 × 10_−4_). There was a significant difference between our conditions, both in reaction time and accuracy (reaction time: F_2.70,99.72_ = 11.7, pHF = 3.0 × 10_−6_, accuracy: F_2.63,97.16_ = 3.8, pHF = 0.016). Additionally, learning was revealed by an interaction between condition and time, again in both measures (reaction time: F_3,111_ = 12.3, p = 5.3 × 10_−7_, accuracy: F_3,111_ = 4.1, p = 0.009).

During the second half of the experiment, after learning has occurred, there was a clear benefit for the Old-ro displays compared to the new displays (Figure 2B/2C). Participants were faster (Old-ro: 1.16 ± 0.44 s, New: 1.28 ±0.46 s, t_37_ = 6.4, p = 1.9 × 10_−7_, d = 0.98) and more accurate (Old-ro: 91.34 ± 8.51%, New: 88.70 ± 9.80%, t_37_ = −3.7, p = 7.0 × 10_−4_, d = −0.10 BF_10_ = 41.81) in locating the target. Bayesian analysis of these results reveal very strong evidence for this benefit in accuracy (BF_10_ = 41.81) and extreme evidence in reaction time (BF_10_ = 8.2 × 104). We thus unequivocally find the classical contextual cueing effect.

Additionally, when we compare the effect of a predictive target location (Targ-so) to no predictive context at all (New) we find no differences in accuracy (Targ-so: 89.50 ± 9.80%, New: 88.70 ± 9.80 %, t_37_ = −1.4, p = 0.17). However, we do find a marginal effect in reaction time (Targ-so: 1.24 ± 0.45ms, New: 1.28 ± 0.46 ms, t_37_ = 2.0, p = 0.05). A Bayesian analysis indicated inconclusive evidence for the presence of an effect on reaction time (BF_10_ = 1.02) and anecdotal evidence for the null hypothesis of no effect for accuracy (BF_10_ = 0.43). Considering we do find a strong contextual cueing effect, this indicates that the effect of spatial predictive context far exceeds any effect of temporal predictive context, when both operate in isolation. Participants were indeed markedly faster finding the target in scenes with spatial predictive context compared to when only target locations were predictive (Old-ro: 1.16 ± 0.44 ms, Targ-so: 1.24 ± 0.45ms, t_37_ = − 3.9, p = 3.6 × 10_−4_, d = −1.03). The same was true for accuracy (Old-ro: 91.34 ± 8.51%, Targ-so: 89.50 ± 9.80%, t_37_ = 2.8, p = 8.0 × 10_−3_, d = 0.20). A Bayesian analysis clearly favors the alternative hypothesis (BF_10_ = 77.00 for reaction times and BF_10_ = 5.00 for accuracy).

Importantly, testing the key hypothesis of our experiment, we find no benefit of presenting scenes in a predictable order. Participants were on average equally fast in finding the target in both type of Old displays (Old-ro: 1.16 ± 0.44 ms, Old-so: 1.15 ± 0.44 ms, t37 = 0.7, p = 0.50). We find similar results for accuracy (Old-ro: 91.34 ± 8.51%, Old-so: 91.83 ± 9.01%, t_37_ = −0.6, p=0.56,). The Bayesian analysis of these results (BF_10_ = 0.22 for reaction time and BF_10_ = 0.21 for accuracy) indicate that the data are 4 – 5 times more likely under the null than under the alternative hypothesis. This means participants did not benefit from adding temporal predictive context to spatial predictive context. From this we can conclude that there is no effect of temporal predictive context in our experiment.

It is possible that the tendency to benefit from temporal or spatial predictive context is governed by individual differences. Perhaps people who are sensitive to spatial predictive context, will not pick up on temporal predictive context, and vice versa. An argument can also be made for the opposite: only people who show sensitivity to spatial predictive context are likely to be sensitive to temporal predictive context. To explore these possibilities, we divided participants into two groups: those with no sensitivity to spatial predictive context (i.e., those who lacked a contextual cueing effect in general, as defined by a RT benefit for Old displays (Old-ro/Old-so combined) compared to New) and those who did show a contextual cueing effect. We then tested these groups separately to see if there is a difference within the Old conditions (Old-ro versus Old-so) for either group. However, there were again no differences in the speed with which the target was found between Old-ro and Old-so displays in both the group of participants who did show contextual cueing (n = 24, Old-ro: 1.11 ± 0.17 ms, Old-so: 1.11 ± 0.18 ms, t23 = 0.52, p = 0.61) and the group of those who did not (n = 14, Old-ro: 1.28 ± 0.15 ms, Old-so: 1.24 ± 0.12 ms, t13 = 0.44, p = 0.67) (Figure 3). Bayesian analysis revealed moderate evidence for the Null hypothesis of no difference in both the Contextual Cueing group (BF10 = 0.24) and the No Contextual Cueing group (BF_10_ = 0.29).

**Figure 3.**
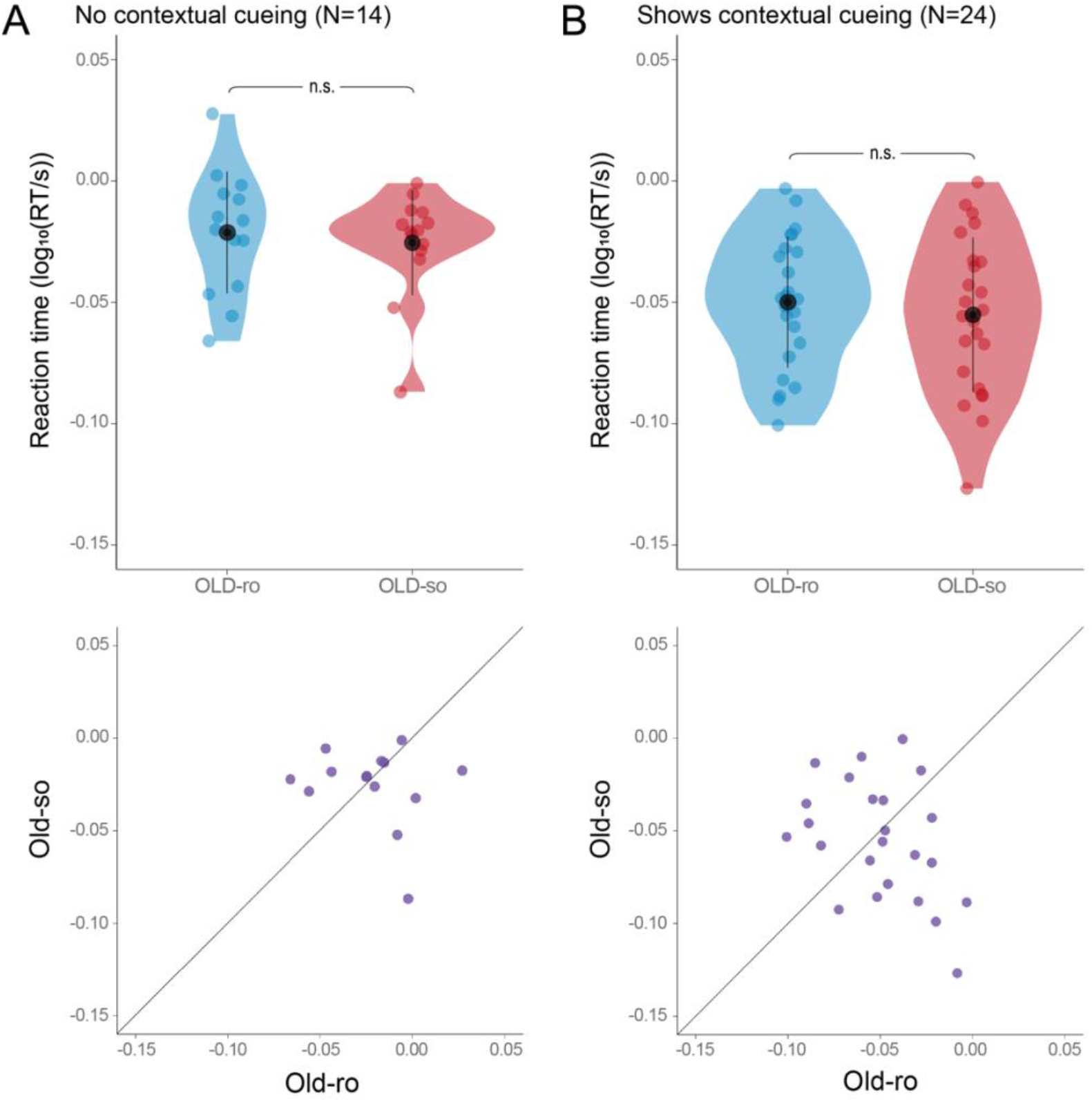
Individual differences in learning of OLD scenes. Reaction time in the Old-ro and Old-so conditions of the second half of the experiment (block 19-36) for participants who revealed no difference between New and Old conditions (A) and participants who were significantly faster on Old compared to New trials (B). Dots in all panels represent individual participants.

**Figure 4.**
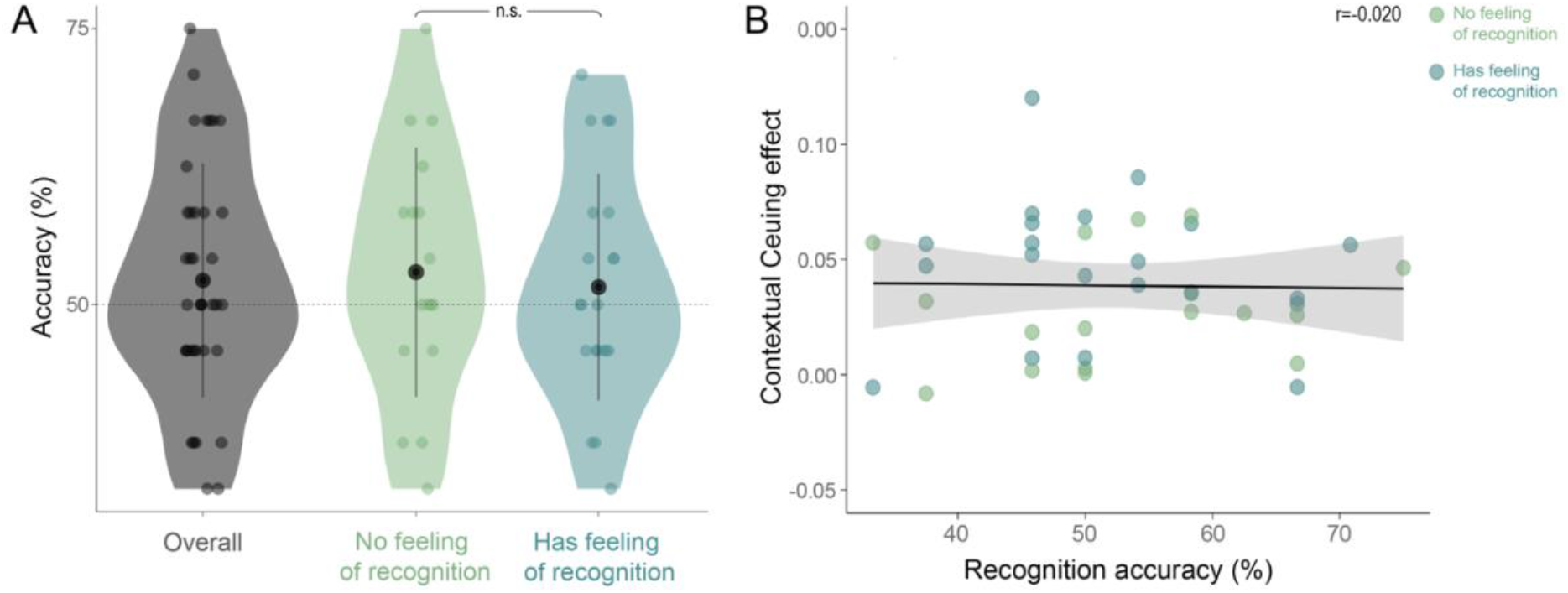
Results recognition task 1. (A) Accuracy in recognizing search displays as either Old or New for the whole sample (left, grey), and split by whether participants indicated a feeling of repeated displays during the main task (right, blue) or not (middle, green). (B) Contextual cueing effect during second half of the main task (New minus Old trials) for the same groups as in (A), as a function of recognition performance. Dots in both panels represent individual participants.

**Figure 5.**
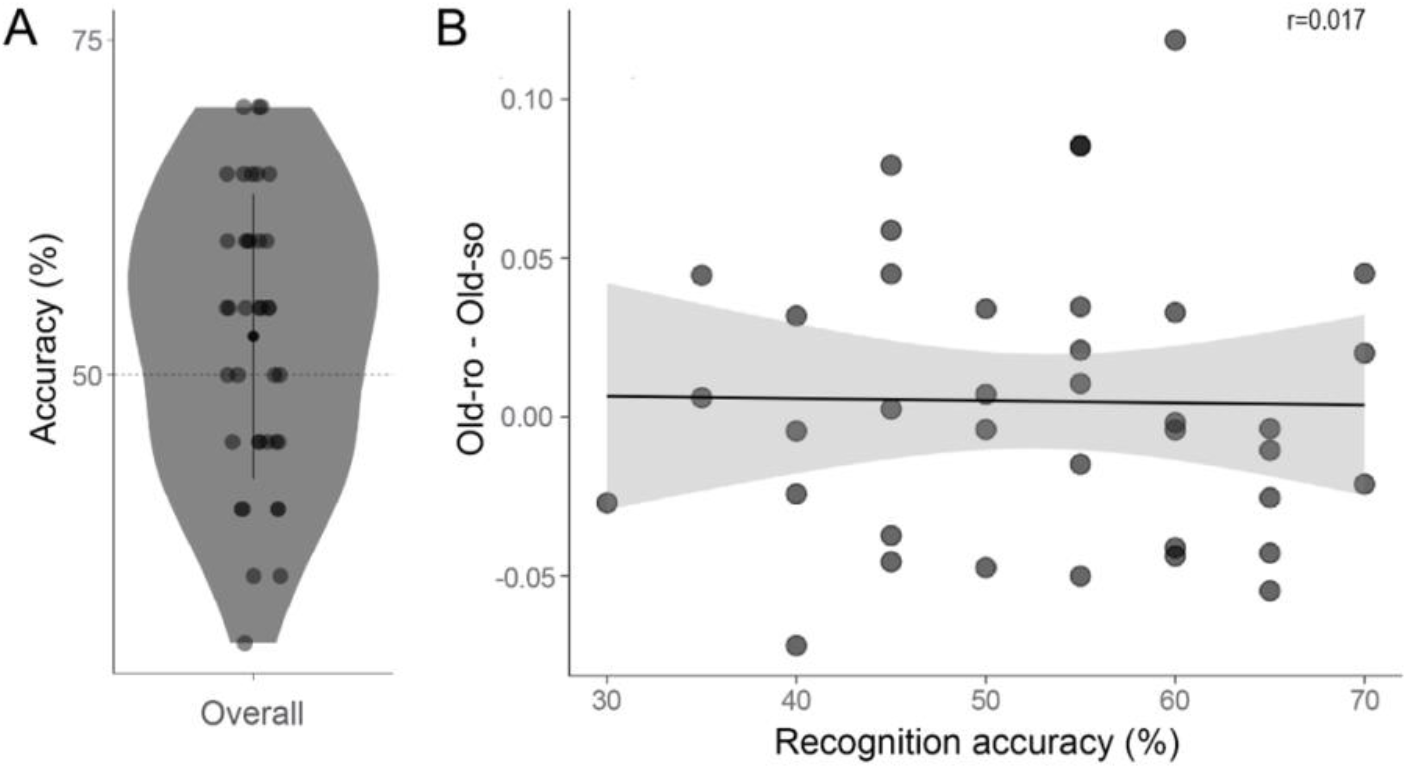
Results recognition task 2 (A) Accuracy in recognizing the order of 2 search displays as either correct or incorrect, for the whole sample. (B) The effect of adding temporal to spatial predictive context task (defined as RT difference between Old-ro and Old-so trials) during second half of the main task as a function of recognition performance. Dots in both panels represent individual participants.

Taken together, these results demonstrate that people did not make use of temporal predictive context in addition to spatial predictive context during our visual search task. This lack of sensitivity to temporal predictive context is independent from individual sensitivity to spatial predictive context.

### Recognition tasks

#### Task 1

Participants were at chance level in correctly identifying displays as Old or New (52.19 ± 10.60%, t-test versus chance level t_37_ = 1.27, p = 0.21, BF_10_ = 0.37). When probed before the recognition task, over half (21/38) of our participants reported to have a feeling that some displays were shown multiple times over the course of the experiment. However, these participants were not more accurate compared to participants who did not report such subjective feelings of recognition (Feeling of recognition: 51.59 ± 10.25%, No feeling of recognition: 52.94 ± 11.29%, t_32.784_ = 0.38, p = 0.70, BF_10_ = 0.34). This is indicative of participants’ subjective report of recognition not reflecting actual recognition performance. Neither of the two groups differed from chance (Feeling of recognition t_20_ = 0.71, p = 0.49, BF_10_ = 0.29, No feeling of recognition: 52.31%, t_16_ = 1.07, p = 0.30, BF_10_ =0.41); we thus found no evidence of explicit memory for the Old displays. Additionally, we did not find a relationship between recognition accuracy and the contextual cueing effect (as defined by the difference between New and Old conditions, r = −0.02, t_36_ = −0.12, p = 0.91, BF_10_ = 0.36).

#### Task 2

Participants were at chance level for recognizing the order of two displays as correct or incorrect (52.89 ± 10.63%, t-test versus chance level t_37_ = 1.68, p = 0.10, BF_10_ = 0.63). A limited number of participants (4/38) reported to have a feeling that the order of some displays was repeated during the experiment. We therefore did not analyze these groups separately. Participants who were better at recognizing the order of displays, did not show a stronger effect of adding temporal to spatial predictive context: we did not find a relationship between recognition accuracy and the difference between random and structured order (r = −0.0167, t_36_ = −0.10, p = 0.9208, BF_10_ = 0.36).

## Discussion

The aim of this study was to investigate the potential benefit of temporal predictive context over and above spatial predictive context during visual search. More specifically, we used a contextual cueing paradigm to investigate whether people can exploit temporal predictive context in addition to spatial predictive context. We found a clear and strong benefit for old displays compared to novel displays, confirming the strong influence of spatial predictive context during visual search. However, we did not observe an effect of repeating old scenes in a consistent order compared to presenting them in a random order. This was also true when we separated participants who were sensitive to spatial predictive context from those who were not. We thus found evidence that temporal predictive context is not exploited when locating a target in a complex scene using a typical visual search task. We discuss these findings below.

As anticipated, we were able to replicate the contextual cueing effect. This phenomenon is strong and robust and seen as evidence of spatial predictive context guiding selective attention through complex scenes (7,22). Moreover, we found evidence neither for explicit knowledge of old scenes, nor for explicit knowledge of the order in which they were presented, as participants performed at chance level for all recognition tasks. This is thought to be evidence that contextual cueing is an entirely implicit effect (5,7,11). Though this finding is in line with previous research, the support for the null-hypothesis of chance-level performance, as expressed by a Bayes factor, was anecdotal. Interestingly, Spaak & de Lange (11) found that participants who revealed *more* explicit knowledge of repeated scenes had a *weaker* contextual cueing effect. We were, however, not able to replicate this negative relationship in the present study. An important difference between the two studies is that we had fewer old scenes: 12 instead of 20. Therefore, we might have been under-powered to detect above chance performance with this relatively small sample of displays (23). In the Spaak & de Lange study participants who reported to have some subjective awareness of scenes being repeated, were better (and above-chance) at recognizing old scenes. Although over half of our sample reported such a subjective feeling of awareness, these participants were not more accurate in recognizing old scenes, rendering this subjective report unreliable in our sample.

Our main finding is that temporal predictive context was not exploited during our visual search task. We further explored this finding by investigating participants who were sensitive to spatial predictive context separately from those who were not. We speculated that if there is a trade-off at the level of individual differences, it might be that participants who are *less* sensitive to spatial predictive context, might be *more* sensitive to temporal predictive context. It is important to acknowledge that visual search is an inherently spatial task. Even in the temporal predictive context conditions, spatial location(s) predicted spatial location(s) on the next trial. From this, it follows that in order to even detect temporal context, spatial context needs to be processed first. Therefore, an alternative possibility is that only participants who are sensitive to spatial predictive context, are also sensitive to temporal predictive context. This is, however, not what we found. Instead, we found no effect of temporal predictive context, whether participants were sensitive to spatial predictive context or not.

We found a marginal effect when only target location predicted the next target location (Targ-so). We included this condition to be able to address the question to what extent a possible effect of temporal predictive context elicited by the entire scene (Old-so) might be due solely to the repetition of the target, independent of the rest of the spatial configuration. The Targ-so condition had a limited set of target locations compared to the completely novel displays, which in turn would elicit a small amount of spatial predictive context. This might have resulted in the marginal benefit that we found.

It might be that if spatial context is fully predictive of the target location, temporal predictive context can no longer contribute in a significant way. This is in line with the notion of a stronger cue blocking the association with a weaker cue (24,25). It would be interesting to see whether temporal predictive context is taken into account when the spatial predictive context is more uncertain, for instance by changing the location of a subset of the distractors, which attenuates but does not abolish the contextual cueing effect (11,24,25). This might additionally explain why our null finding is at odds with the phenomenon of intertrial contextual cueing (26,27). This effect is a search benefit for finding a target on trial N, with the predictive context being an entire display (distractors & target location) on trial N-1. Ono et al. concluded that the “visual system is sensitive to all kinds of statistical consistency and will make use of predictive information whether it is presented in a single trial or across trials.” In our case, however, the visual system seems to ignore the temporal consistency across trials. An important distinction to be made is that with intertrial contextual cueing, spatial predictive context is missing on trial N. The only context predicting target location on trial N is therefore the spatial context on trial N-1. If spatial predictive context is dominant, this might explain why temporal predictive context *can* be exploited in intertrial contextual cueing: there is no spatial predictive context on trial N ‘hindering’ the process. Given our results, we predict that adding spatial predictive context on trial N in an intertrial contextual cueing experiment abolishes the effect. Supporting this notion, Olson & Chun (28) found that exposing participants to sequences of displays with multiple distractors, and a target at the end of the sequence, does yield a benefit in detecting the target when these sequences are structured compared to random. Future research is needed to establish this.

Finally, it could be that we did not use enough repetitions for participants to pick up temporal predictive context. We repeated the displays 36 times, which is plenty for the contextual cueing effect to appear, but we cannot exclude the possibility that more learning is required to pick up on temporal regularity. Research investigating temporal statistical regularities commonly uses equally many or even fewer repetitions (28–30), but an important difference might be that stimuli typically used are easily processed, such as objects in isolation. Visual search scenes are more complex, representing multiple target-distractor relations. Even though recent work shows there is also a component of scene memory to contextual cueing (31,32), this might not be strong enough to enable one ‘scene’ to function as a predictor of the next, analogous to how an object predicts the next one in typical temporal statistical learning tasks. A previous study exposed observers to sequenced information in addition to, but independent of, spatial predictive context. Signs of sequence learning were visible after 96 repetitions (33). It is possible that this many repetitions of the spatial context in our study would have allowed exploitation of temporal predictive context. We question, however, whether learning will then remain implicit, introducing another source of variance.

Our results have implications for how we think about the role of predictive context in visual perception. Though both spatial and temporal context are of importance for visual perception (34), our findings imply that extracting and using context might not be as automatic as previously assumed. In our experiment we find that spatial predictive context is favored. This might be due to a modality-specific bias, leading to a dominance of spatial predictive context in visual perception (35–39). However, sensitivity to one type of context over the other might also be task-dependent.

It is possible that temporal predictive context is encoded by observers, but not exploited. Within contextual cueing it has been shown that the exploitation of spatial predictive context knowledge depends on attention, even though its acquisition does not (16,24). Additionally, learned predictive context can be visible at the neural level without being expressed in behavior (13,15). Therefore, not finding an effect of temporal order in behavior in our task does not necessarily imply that this predictive context is not encoded.

Concluding, in our study we found evidence that observers do not benefit from temporal context on top of spatial predictive context during visual search. We argue that since people are focused on locating an object in space, spatial context is readily encoded and subsequently exploited, while temporal context may not be. More research is needed to investigate the specific conditions that would allow for temporal contingencies to have an effect on search behavior, and to ultimately understand how spatial and temporal predictability jointly shape our perception.

## Acknowledgements

We would like to thank Izana Bayerman for her help with collecting the data.

